# Cross-domain interactions induce community stability to benthic biofilms in proglacial streams

**DOI:** 10.1101/2023.01.31.526486

**Authors:** Susheel Bhanu Busi, Hannes Peter, Jade Brandani, Tyler J. Kohler, Stilianos Fodelianakis, Paraskevi Pramateftaki, Massimo Bourquin, Leïla Ezzat, Grégoire Michoud, Stuart Lane, Paul Wilmes, Tom J. Battin

## Abstract

Cross-domain interactions are an integral part of the success of complex biofilms in natural environments. Here, we report on cross-domain interactions in biofilms of streams draining proglacial floodplains in the Swiss Alps. These streams, as a consequence of the retreat of glaciers, are characterized by multiple environmental gradients and stability that depend on the time since deglaciation. We estimate co-occurrence of prokaryotic and eukaryotic communities along this gradient and show that key community members have disproportionate effects on the stability of co-occurrence networks. The topology of the networks was similar independent of environmental gradients and stability. However, network stability was higher in the streams draining proglacial terrain that was more recently deglaciated. We find that both pro- and eukaryotes are central to the stability of these networks, which fragment upon the removal of both pro- and eukaryotic taxa. These ‘keyplayers’ are not always abundant, suggesting an underlying functional component to their contributions. Thus, we show that there is a key role played by individual taxa in determining microbial community stability of glacier-fed streams.

## Introduction

Biofilms represent the dominant microbial lifestyle in streams and rivers^1^. There, these matrix-enclosed microbial communities colonise sediment surfaces and can regulate critical ecosystem processes^1^. Stream biofilm communities are highly diverse, harbouring members of all domains of life, including viruses. This biodiversity fosters biotic interactions, such as those between algae and bacterial heterotrophs, which contribute to the stability of ecological communities^2^. Given the multitude of interacting taxa and the small spatial scales at which interactions occur, their direct observation is, however, not possible. Instead, patterns of taxa co-occurrence across samples can be used to infer microbial interactions. These co-occurrence patterns are often usefully represented as ecological networks, which allow us to explore emergent properties, such as the density of interactions, clusters of interacting taxa or the stability of networks against fragmentation. For example, studying bacterial co-occurrences across a dendritic stream network, Widder *et al*. found evidence for the role of spatial and hydrological processes in shaping co-occurrence network structure and stability^3^.

Overall, the environment of proglacial streams is extreme. Low water temperature coupled to high turbidity and oligotrophy as well as snow- and ice-cover over extended times collectively contribute to rendering these environments extreme. Highly unstable stream channels further contribute to these extreme conditions, making it difficult for benthic biofilms to establish^4^. This is particularly true for glacier-fed streams (GFS) that develop into braided channels, and which commonly are dynamic with channel changes on a diel basis. Further downstream, these effects become alleviated notably by biogeomorphic succession as plant communities begin to exert substantial resistance to lateral channel erosion^5^. GFS channels start to consolidate, thereby further increasing the habitability of the GFS ecosystem. Towards the edge of the proglacial floodplain, tributaries (TRIB) fed by groundwater and snowmelt drain terrasses that are slightly elevated and often disconnected from the meltwaters in the GFS channels^6^. The environment in TRIB is generally more stable than in GFS^4^, which is reflected by the microbial communities in these streams^7,8^. In fact, despite their close spatial proximity, GFS and TRIB host biofilms that differ in terms of biomass, composition, and diversity^7,8^. GFS will become increasingly fed by groundwater and snowmelt as glaciers shrink^9^.

Here, we investigated the properties of cross-domain microbial co-occurrence networks in benthic biofilms in GFS and TRIB within zones with different deglaciation histories in two proglacial floodplains in the Swiss Alps. We hypothesised that the apparent stability of co-occurrence networks in GFS and TRIB changes along downstream and lateral gradients of deglaciation histories and hence environmental stability. To address this, we assessed the stability of cross-domain co-occurrence networks upon removal of keyplayer taxa. Keyplayers are taxa with a central role in maintaining network structure and have been identified in other ecological networks^10,11^. However, the role of keyplayers for structuring communities is unknown. We investigated the variance in bacterial community composition that can be explained by eukaryotic and prokaryotic keyplayers and contrasted this to the variance that can be explained by environmental differences among sites. Our findings highlight the importance of cross-domain interactions for the success of biofilms in proglacial streams.

## Materials and methods

### Sample collection

Benthic sediment from various stream reaches within the Otemma Glacier (Otemma; 45° 56’ 08.4” N 7° 24’ 55.1” E) and Val Roseg Glacier (Val Roseg; 46° 24’ 21.1” N, 9° 51’ 55.1” E) floodplains were collected from the glacier snout to the floodplain’s outlet. In each reach, we collected sandy sediments (0.25 - 3.15 mm) from the benthic zone (0 - 5 cm depth) with flame-sterilised sieves and spatulas. Samples were collected during early (June/July) and late (August/September) summer^8^ and as shown previously by Brandani *et al*. ^8^, the two sample periods did not show differences in terms of community composition and structure. Study reaches were categorised into GFS or TRIB depending on their connectivity to glacier runoff based on visual field observations, drone-based imagery, and physicochemical characteristics^8^. Overall, a total of 136 samples (GFS: 50; TRIB: 86) were collected across both floodplains. These included 68 samples each for the Otemma Glacier and Val Roseg Glacier floodplain, where the exact breakdown of these samples into GFS and TRIB, UP and DOWN (see *Methods*) are listed in Supplementary Table 1.

### Deglaciation histories

We identified past glacier extents from historic orthophotos and maps using SWISSIMAGE journey through time^12^, and the GLIMS glacier inventory^13^. These extents were compared with GLAMOS^14^ frontal variation measurements to verify glacial readvances. Year of latest glaciation was thus interpolated for each sample site, which provided the longitudinal deglaciation history. We further split the reaches of the floodplain into those which were already deglaciated in 2000 (DOWN) and those still glaciated in 2000 (UP) (Supp. Fig. 1a). The lateral gradient is given by the TRIB that drain the terraces on the margins of the proglacial floodplains.

**Figure 1.**
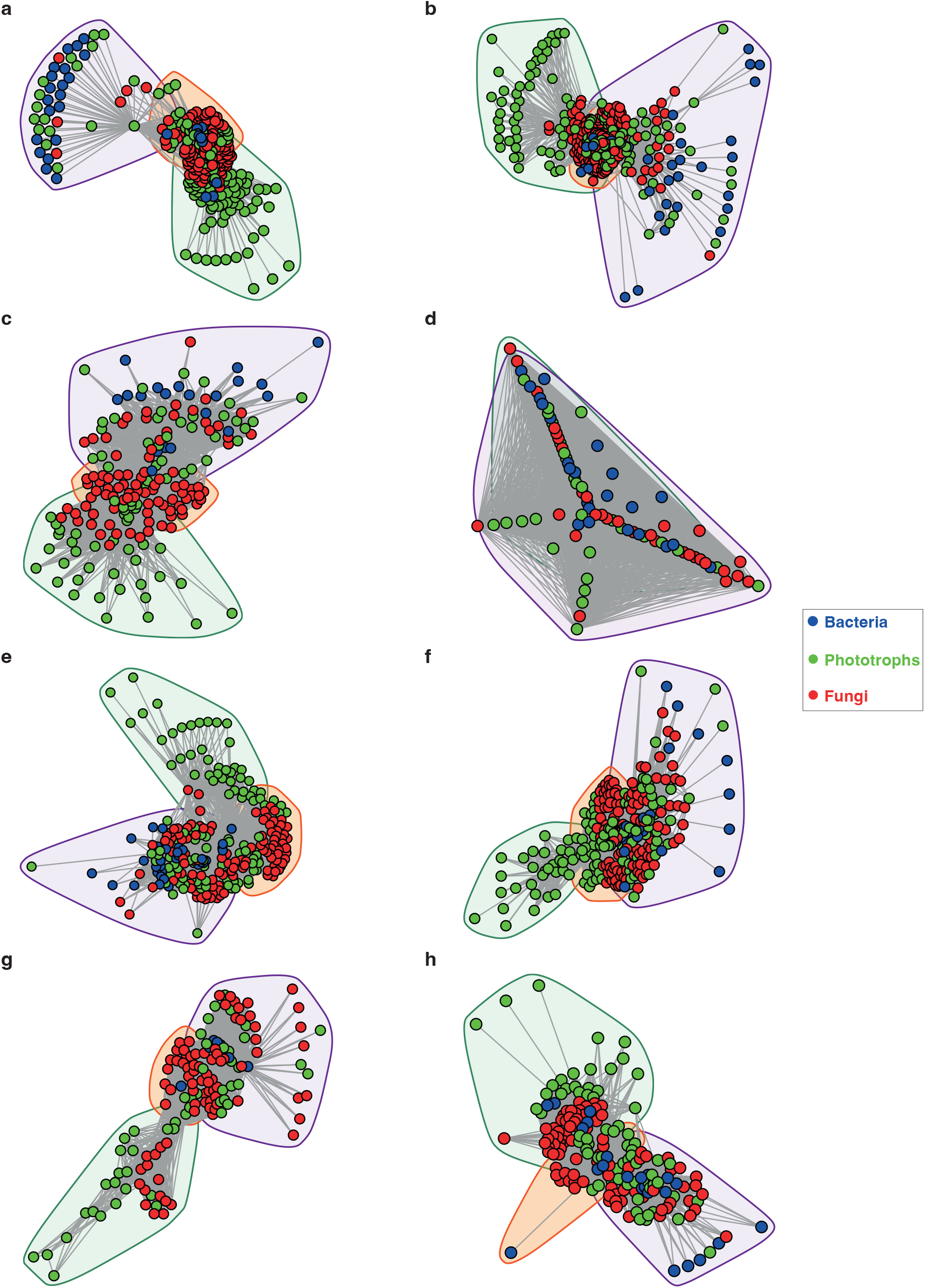
Network structure of Glacier-fed streams and tributary streams. The overall structure of the cross-domain networks from the GFS and TRIB are depicted. (a) GFS from the UP reaches at Otemma, (b) GFS from the DOWN reaches at Otemma, (c) TRIB from the DOWN reaches at Otemma, (d) TRIB from the DOWN reaches at Otemma. From the Val Roseg glacier, the network structures are depicted as follows: (e) GFS from the UP reaches, (f) GFS from the DOWN reaches, (g) TRIB from the UP reaches, (h) TRIB from the DOWN reaches. Each node represents a single amplicon sequence variant (ASV), and the lines represent the edges between them, while the colours indicate bacteria, phototrophs and fungi. The convex hulls indicate clusters identified based on Louvain clustering of the overall network.

### Benthic algal biomass

Benthic algal biomass was estimated as chlorophyll *ɑ* using a modified ethanol extraction protocol ^15^. For this, the sediment (ca. 2 g) samples were treated with 5 ml of 90 % EtOH and then placed in a hot water bath (78 °C, 10 min), followed by an incubation in the dark (4 °C, 24 h). They were thereafter vortexed, centrifuged, and the supernatant read on a plate reader at 436/680 nm (excitation/emission). Chlorophyll *ɑ* concentrations were inferred from a spinach standard and normalised by the sediment dry mass (DM).

### Metabarcoding library preparation, and sequencing

A previously established protocol^16^ utilising phenol-chloroform was used for DNA extraction from benthic sediments (ca. 0.5 g). After initial processing, the DNA samples were diluted to a final concentration of ≤ 2-3 ng/ul. For the 16S rRNA gene metabarcoding analyses, we used the methodology previously described in Fodelianakis et al.^17^, where the V3-V4 hypervariable region of the 16S rRNA gene were targetted with the 341F/785R primers. This was done in line with the 16S library preparation Illumina guidelines for the MiSeq system. The eukaryotic 18S rRNA gene metabarcoding library preparation was performed similarly but using the TAReuk454F-TAReukREV3 primers^18^. Based on the MiSeq manufacturer’s protocol, amplicon libraries were prepared where a second PCR was used to add dual indices to the purified amplicon PCR products. This allowed for extensive multiplexing of samples on a single sequencing lane of the MiSeq (Illumina) platform after quantification and normalisation. Samples were subsequently sequenced using a 300-base paired-end protocol in the Lausanne Genomic Technologies Facility (Switzerland).

### Metabarcoding analyses

For the 16S and 18S rRNA metabarcoding data analyses, a combination of Trimmomatic v0.36^19^ and QIIME2 v.2020.8^20^ were used with the latest SILVA database^21^ v138.1 for taxonomic classification of the gene amplicons, i.e. 16S rRNA and 18S rRNA. From the 16S rRNA amplicon dataset, non-bacterial amplicon sequence variants (ASVs), i.e., archaea, chloroplasts, and mitochondria, were removed from all downstream analyses. The dataset was not rarefied for the analyses. The rationale behind discarding the archaeal reads was that the primers used were not designed, and are therefore not optimal, for detecting all lineages of archaea^22^. A total of 192 sample libraries were generated for the 16S rRNA sequencing and paired-end sequencing produced a total of 15,140,043 reads, with an average of 89,586 reads per sample. However, only 136 were included in the analysis due to absence of paired 18S rRNA information for 56 samples. Meanwhile, singletons and ASVs observed only once were discarded. For the 18S rRNA amplicon dataset, 136 amplicon sequence libraries from sediment samples were generated (17 samples were discarded due to DNA extraction and amplification issues). The paired-end sequencing generated a total of 10,837,518 reads, with an average of 64,127 reads per samples. The 18S ASVs were further clustered into operational taxonomic units (OTUs) based on a 97% identity threshold using the de novo clustering method in *vsearch*, which has been implemented in QIIME2. Non-phototrophic eukaryotes except fungi and protists were discarded from the 18S rRNA amplicon dataset in all downstream analyses. The 18S rRNA dataset was also not rarefied and any singletons/OTUs observed in only one sample were removed from downstream analyses, resulting in an 18S rRNA phototrophs and fungi dataset of 429 OTUs.

### Co-occurrence networks

To study potential interactions between pro- and eukaryotes, co-occurrence network analyses were performed with samples meeting specific criteria. These included: 1) the presence of both 16S and 18S sequence data for each sample, and 2) samples had to be categorised the same way across both samplings to ensure replicability (i.e., either designated as GFS or TRIB for both samplings as described by Brandani *et al*.^*8*^). Due to the dynamic nature of proglacial streams, GFS tend to migrate, leaving some sites dry or under the influence of TRIB, or even flood samples previously under the influence of TRIB streams. Hence, this approach was adopted to avoid potential confounders arising from miscategorised streams. Subsequently, to reduce the noise and overall computational effort, any ASVs found in less than 5% of the samples were discarded from the 16S dataset for the co-occurrence networks. Co-occurrence networks between 16S and 18S (i.e., phototrophs and fungi) were constructed using an average of the distance matrices created from SparCC^23^, Spearman’s correlation^24^, and SpiecEasi^25^ where the networks were constructed using the Meinshausen and Bühlmann (mb) method (Meinshausen and Bühlmann, 2006). Networks were constructed across reaches, for UP and DOWN segments separately, and across GFS and TRIB for both Otemma and Val Roseg floodplains. Since our analyses are based on amplicon sequence data alone, we focused on the positive interactions across domains to assess potential mutualism within the microbiome. While reports suggest that negative interactions are indicative of co-exclusion mechanisms, especially in human microbiomes^26^, the paucity of information available, especially in poorly characterised ecosystems may be insufficient to establish via amplicon sequencing data.

To detect communities in the network analyses, we used the Louvain clustering algorithm^27^, removing clusters with less than 5 nodes. Herein, each community is defined as nodes within the graph with a higher probability of being connected to each other than to the rest of the network. Following this, we calculated network topology measures, including nodes and edges number, number of clusters, diameter, edge-density, and modularity. The correlation matrix was visualised using the *igraph* package^28^ in R v4.0.3^29^. Centrality measures, degree and betweenness, were also estimated per node, using the *igraph* v1.3.4 package. The fragmentation (*f*) of the network was determined as the percentage of the number of disconnected subgraphs over the overall nodes in each network^3^. Fragmentation was estimated iteratively by the removal of each keyplayer, i.e., top 10 nodes with both a high degree and a high betweenness in each graph. This information was further used for the subsequent generation of network topologies such as the number of clusters following initial Louvain clustering of the network.

### Community analyses

To explore the role of keyplayers in structuring biofilm communities, we used constrained ordinations (db-RDA, R function vegan:capscale) using Bray-Curtis distances. We employed a forward selection strategy (vegan:ordistep) to identify a non-redundant and significant (p<0.01) set of both pro- and eukaryotic keyplayers that explained variance in the bacterial community. We performed this analysis on each floodplain individually. Prior to db-RDA, wisconsin-double standardisation was applied to the bacterial community. The relative abundances of keyplayers were then provided as constraints for stepwise model creation (using 199 permutations). Model significance was evaluated for each RDA axis and explained variance of the constraints was extracted. To contrast variance in bacterial community composition that could be explained by keyplayers, we performed a similar analysis using environmental parameters. For this, important environmental parameters including pH, water temperature, specific conductivity, dissolved oxygen (DO), turbidity and major ions and nutrients were first standardised and then supplied to forward selection in db-RDA as described above.

### Data Analysis

All statistical analyses were performed in R v4.0.3. The *ggplot2*^*30*^ package was used for generating plots in R, while *patchwork* (https://github.com/thomasp85/patchwork) and Adobe Illustrator were used to arrange the figures as displayed.

## Results

### Cross-domain interactions underlie stream community structure in proglacial floodplains

In both proglacial floodplains, GFS and TRIB harbour diverse microbial communities including bacteria, fungi and phototrophic eukaryotes (Supp. Fig. 1). Based on covariation of taxon abundances across samples, we built co-occurrence networks. These networks were based on 1,090 nodes including both pro- and eukaryotes, with an average of 61,115 edges. The topological characteristics of the individual networks yielded similar metrics, such as density, modularity, assortativity and transitivity (Supplementary Table 1). In all networks, except OtemmaDOWN, we identified three dense clusters of co-occurring taxa, one with a majority of phototrophs, another comprising mainly prokaryotes, and an intersecting third cluster composed of microbial eukaryotes including fungi and prokaryotes.

Next, we assessed the relative abundance of taxa present in the networks at the family level. Across both floodplains and stream types, we found that Acetobacteraceae were significantly overrepresented in networks constructed from UP compared to DOWN reaches (two-way ANOVA, adj. *p* < 0.05, Supp. Fig. 2a-b and 3b). On the other hand, Comamonadaceae were significantly overrepresented in DOWN networks (two-way ANOVA, adj. *p* < 0.05), especially in TRIB (Supp. Fig. 2b and 3b). We also found that Chrysophyceae were overrepresented in UP networks, while Diatomea were decreased in UP networks (two-way ANOVA, adj. *p* < 0.05) (Supp. Fig. 2c-d and 3c-d). Chytridiomycota, parasitic fungi infecting algae ^31^, were prevalent in both GFS and TRIB networks, but their abundance did not significantly differ across UP or DOWN sites. However, Zoopagomycota, also parasitic fungi^32^, were considerably enriched in DOWN reaches across stream types and floodplains (Supp. Fig. 2e-f and 3e-f; adj. *p* < 0.05, Two-way ANOVA).

**Figure 2.**
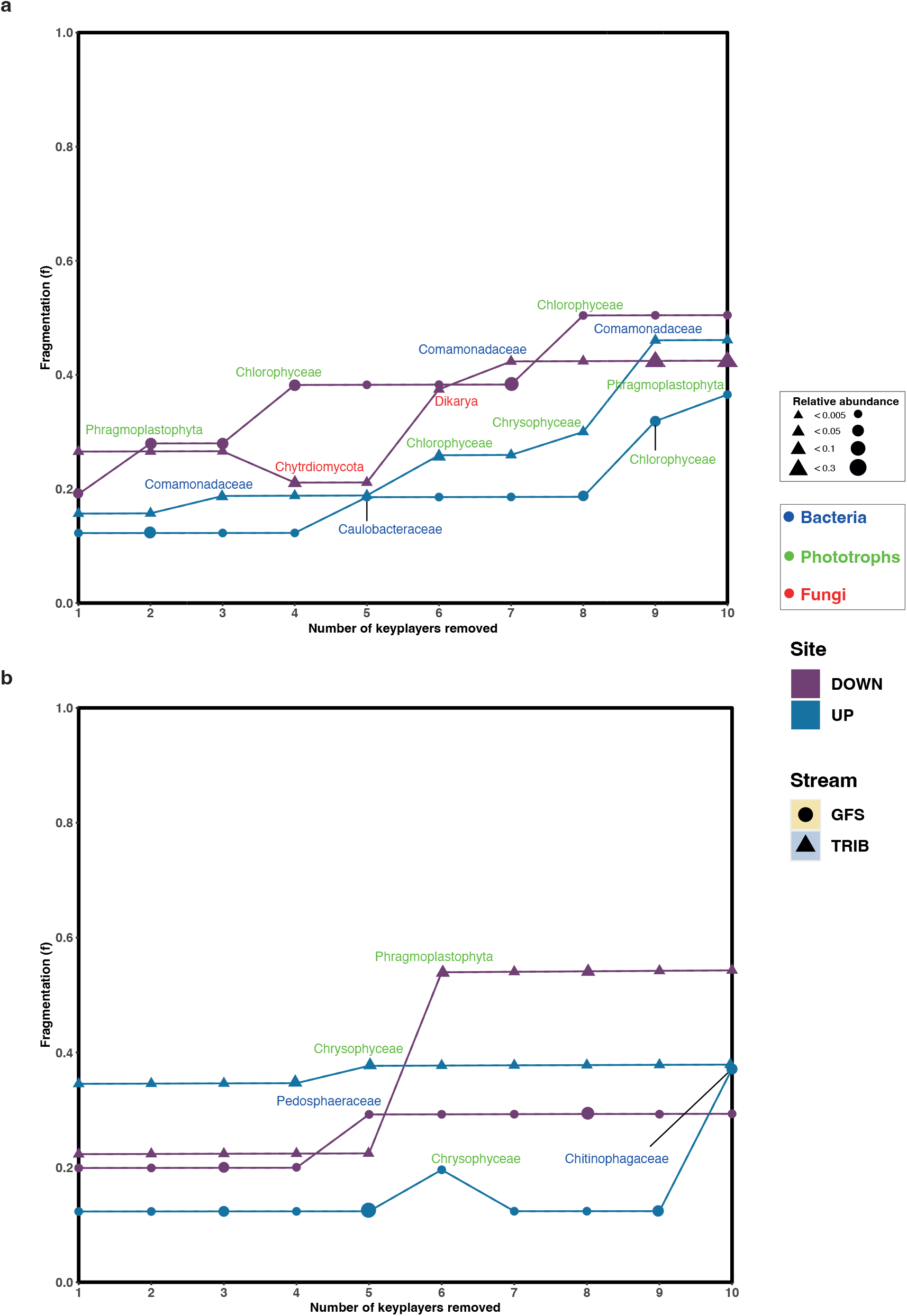
Keyplayer removal leads to fragmentation of the network. The change in fragmentation (*f*) for (a) Otemma and (b) Val Roseg are indicated in the line plots, where *f* was recalculated after each keyplayer was removed from the network. The size of the symbols indicates the relative abundance of the individual ‘keyplayers’ within the 16S or 18S data respectively.

### Apparent stability of co-occurrence networks

Based on our observations of differential abundance patterns across stream types and deglaciation gradients, we further assessed the contributions of the individual taxa to the overall network. For this, we first identified potential keyplayers within each network by identifying the top 10 nodes with both a high degree and a high betweenness in each network (Supp. Fig. 4 and 5). For example, taxa classified as Dikarya, Phragmoplastophyta, Chlorophyceae, Cryptomycota, and Diatomea, along with an ASV classified as Burkholderiales, were determined to be keyplayers in the GFS network at the UP segment of the Otemma Glacier floodplain (Supp. Fig. 4a). Conversely, at the DOWN segment of the same floodplain, Burkholderiales, Phragmoplastophyta, Xanthophyceae, Chrysophceae, and Dikarya, for instance, were identified as keyplayers. Similarly, in the UP segment of the Val Roseg Glacier floodplain, Dikarya, Phragmoplastophyta, Gemmatales, Burkholderiales, Cryptomycota, and Diatomea, for instance, were identified as keyplayers contributing to the network topology (Supp. Fig. 5a, and 5c). Finally, we found various bacteria (e.g., Rhodobacterales, Sphingomonadales) and fungi (e.g., Chytridiomycota) to be keyplayers in the DOWN reaches within the Val Roseg floodplain (Supp. Fig.5b, and 5d).

To further understand the role of the keyplayers in community structure and their effect on overall network stability, we first assessed network fragmentation upon their removal. For this, the numbers of clusters based on Louvain clustering were determined for each network, following which, a keyplayer was removed. The fragmentation (*f*) of the network was assessed before and after iterative removal of the top ten keyplayers. Interestingly, we found that in the Otemma Glacier floodplain (Fig. 2a), the fragmentation of the networks constructed from the GFS in the DOWN reaches, increased upon removal of two to three keyplayers, while the TRIB fragmentation increased upon removal of five keyplayers. The UP networks, however, appeared more stable, where fragmentation occurred only upon removal of five or eight keyplayers. In Val Roseg, especially in TRIB (Fig. 2b), the overall fragmentation of the microbial network was higher (*f*_mean_=0.48) compared to GFS (*f*_mean_=0.18) upon removal of four or five keyplayers.

Finally, we unravelled the role of keyplayers for biofilm community composition. Constrained ordinations revealed that both, prokaryotic as well as eukaryotic keyplayers can explain a substantial fraction of bacterial community dissimilarity at the floodplain scale (Fig. 3). Specifically, the relative abundance of prokaryotic keyplayers explained 35.0% and 25.4% of variance in bacterial community similarity in Val Roseg and Otemma, respectively. While eukaryotic keyplayers appeared particularly important for explaining network stability, they played a minor role in explaining bacterial community composition (i.e., 8.5% and 2.4% of explained variance in Val Roseg and Otemma, respectively). This is surprising, particularly in relation to the variance in bacterial community composition that could be explained by environmental conditions, which accounted for a mere 16.5% and 14.5%, respectively. The retained environmental parameters, including streamwater temperature, nutrients (i.e., NO_2_, PO_4_) and DOC concentration explain differences among TRIB and GFS bacterial communities.

**Figure 3.**
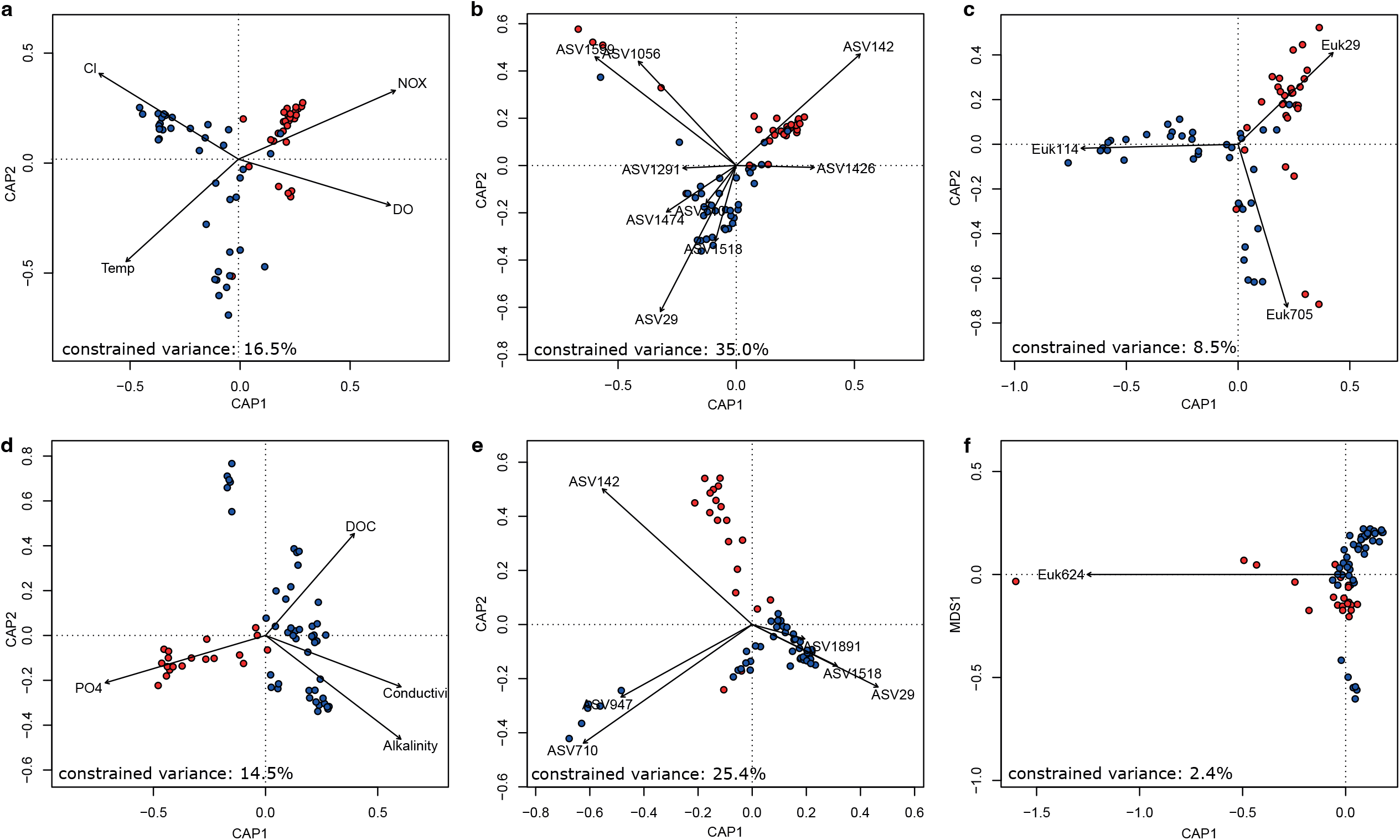
Prokaryotic keyplayers well explains bacterial community composition. Constrained ordination of Val Roseg (a, b, c) and Otemma (d, e, f) floodplain samples revealed that prokaryotic keyplayers (b, e), as identified by their position in co-occurrence networks explained most of the variance in Bray-Curtis distance based bacterial community composition. This outweighed the role of key environmental parameters (a, d) and of eukaryotic keyplayers (c, f).

## Discussion

Biotic interactions are a salient property of microbial communities, with evidence of cross-domain interactions reported from various ecosystems, including freshwaters^33,34^, oceans^35^ and glaciers^36^. To date, such interactions have not been studied in proglacial stream biofilms. Our findings suggest that biotic interactions, as inferred from co-occurrence patterns, play a pivotal role in influencing the apparent stability of stream biofilm communities along deglaciation and environmental gradients in proglacial floodplains. Although previous reports showed structural and functional differences of the biofilm communities dwelling in different stream types within proglacial floodplains^7,8^, we found that the overall network topology was similar between both proglacial floodplains, stream types, and deglaciation gradients. This contrasts our expectation of successional imprints owing to deglaciation on co-occurrence networks. On the one hand, biotic interactions may be established very early on during community succession in streams that drain recently deglaciated terrain. Indeed, our sampling design covered the successional timescale of the past 20 (UP sites) and 80 (DOWN) years and both prokaryotic and eukaryotic communities are likely to assemble much faster. On the other hand, functional redundancies across clades may also contribute to the apparent similarity of cross-domain interaction networks. Functionally redundant taxa may transiently occupy the same position in interaction networks and therefore result in similar network topologies. However, additional work will be necessary to relate network topology, taxa position and stability with functional characteristics to substantiate this notion.

Cross-domain networks have the potential to reveal key associations between microbial taxa^37^. We found that biofilms in GFS and TRIB draining recently deglaciated terrain (i.e., UP sites) had relatively more stable networks. This finding suggests that prokaryotic keyplayers are important for the apparent stability of the cross-domain interaction networks of biofilms dwelling in nascent stream ecosystems. Furthermore, our results reveal that keyplayers are typically not among the most abundant community members, suggesting that low abundance taxa may also play important roles in stabilising microbial networks, corresponding to the notion of keystone species^38^. Our findings agree with observations from recent reports^39–41^ highlighting the role of low-abundance taxa in ecosystem function and structure. For example, de Cena *et al*. recently hypothesised that low-abundance taxa, albeit in the human microbiome, act as keystone species, and might often be more metabolically influential within the community^39^. Similarly, Crump *et al*.^40^ identified microbial keystone species that are central to ecosystem-level metabolic activity.

Work on multi-trophic food webs^42^ and agroecosystems^43^ has demonstrated the fragility of ecological networks towards removal of key nodes. Our fragmentation analysis substantiates the notion of keyplayers and their role for the stability of the cross-domain network. Interestingly, we identified several eukaryotes as keyplayers, underscoring their relevance for biofilm structure and functioning. In GFS in Central Asia, Ren *et al*. ^44^ reported that fungi form integral components of cross-domain interactions networks, forming more clustered networks that are less susceptible to disturbances. As highlighted previously for stream biofilms^45,46^, eukaryotic algae serve as sources of organic matter thereby fuelling phototrophic-heterotrophic interactions. Simultaneously, parasitic fungi also foster the release of organic compounds from algae via the ‘fungal shunt’^31^. The prevalence of parasitic fungi has been noted previously in GFS^47^ and other cryospheric ecosystems^48^; our analyses further point to the importance of interactions among parasitic fungi and their algal host in proglacial stream biofilms. Along these lines, Mo *et al*.^*49*^ recently suggested that interactions of microeukaryotes between them in the Lena River continental shelf were more stable compared to that of the estuary, potentially explained by variability in salinity. In contrast, Liu and Jiang (2020), reported that bacteria-bacteria interactions dominate co-occurrence networks in coastal sea waters of Antarctica^50^ and related this to competitive abilities of prokaryotes.

Taken together, the roles of pro- and eukaryotic keyplayers for ecological networks and their stability may very much be context dependent. We argue that, likely driven by the provisioning of organic matter to heterotrophs, eukaryotic algae and their fungal parasites play central roles in biofilm interaction networks. However, we quantified the relative importance of pro- and eukaryotic keyplayers to overall bacterial community structure and found that the relative abundance of prokaryotic keyplayers could explain much of the bacterial community structure. This points towards a hierarchical structuring of interactions among eukaryotic and prokaryotic biofilm members. While eukaryotic primary producers may directly interact with only some bacterial keyplayers, these bacterial keyplayers themselves interact, likely via the exchange of secondary metabolites, with a much larger number of prokaryotes in the biofilm assemblage. Such a hierarchical organisation of interactions is likely sensitive to changes in taxa at the base (i.e., the algal primary producers) whereas functional redundancies may dampen the impacts of taxa replacement. This is particularly relevant in proglacial streams, where low light availability due to suspended particles and substrate instability typically inhibit algal growth. The current retreat of glaciers weakens these controls with potential effects on stream microbial communities.

## Acknowledgements

We would like to acknowledge Kevin Casellini, Nicola Deluigi, Matteo Roncoroni, for their help sampling in the field. We would also like to thank Martina Schön for her help on collecting and interpolating glacial extent. The Hunting and Fishing Office of the Canton of the Grisons gave permission to fly the drone on the floodplain of the Tschierva Glacier. Funding was provided by the Swiss National Science Foundation grant (CRSII5_180241) to Tom J. Battin, Stuart Lane and Paul Wilmes.

## Data availability

The raw sequence data for both 16S and 18S amplicon sequencing are available on NCBI under the accession ID: PRJNA808857. The metadata associated with the sequence data is also available on NCBI along with the data. The processed ASV abundance tables including the taxonomic affiliations are provided as Supplementary Table 3. Additional data required for figure generation are available at https://doi.org/10.5281/zenodo.7524289.

## Code availability

The code for running the initial network generation and subsequent analyses including figure generation can be found at the following repository: https://doi.org/10.17881/0gdr-7705.

## Tables

**Supplementary table 1. Metadata and network topology**.

Glacier metadata including the glacier from which samples were collected, UP or DOWN reaches, and type of stream, i.e., Glacier-fed (GFS) or tributaries (TRIB), are indicated along with network topology measures. The dashed line (----) indicates the ‘millennium cut’ based on which samples were classified as ‘up’ or ‘down’. The solid lines represent the deglaciated history based on the Glacier Extent Database to determine the date since ‘last glaciation’.

**Supplementary table 2. Chlorophyll-ɑ measurements**.

Levels of chlorophyll-ɑ measured at the site for each sample are listed along with the metadata.

**Supplementary table 3. Pro- and eukaryotic keyplayers**.

Abundance information for all ASVs detected in the prokaryote (16S) and eukaryote (18S) datasets are provided alongside their indication of Keyplayers or otherwise.

## Supplementary figure legends

**Supplementary figure 1.**
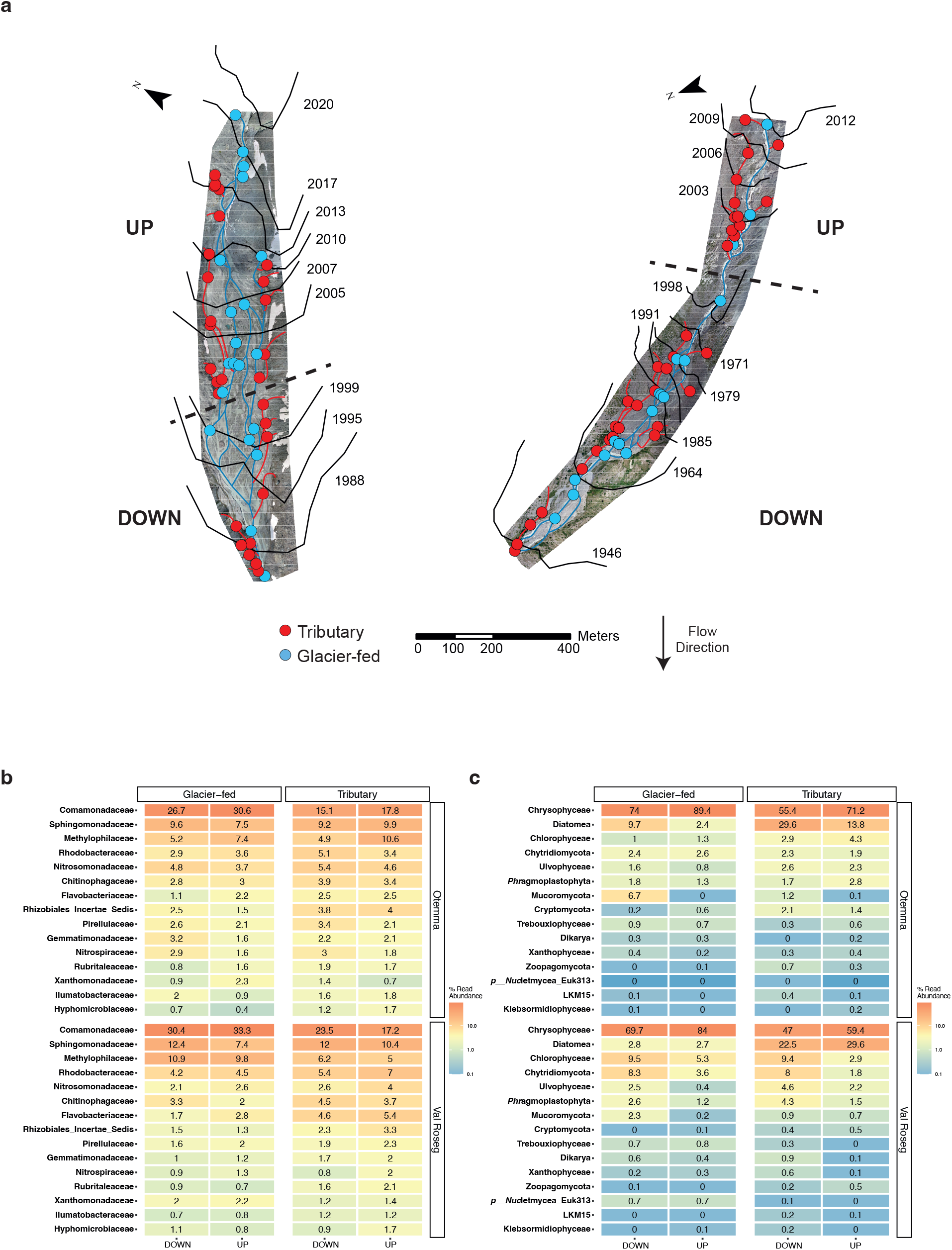
16S and 18S community profiles. (a) Bird’s eye-view of the Otemma (left) and Val Roseg (right) floodplains depicting the mainstem (GFS) and the branching TRIB. The dashed line indicates the ‘Millennium cut’, where samples were classified as ‘up’ or ‘down’ site above and below, respectively. (b) Family-level profiles of the top 15 prokaryotes found in the floodplains across reaches and stream types (GFS and TRIB). (c) Relative abundance of the top 15 eukaryotic taxa.

**Supplementary figure 2.**
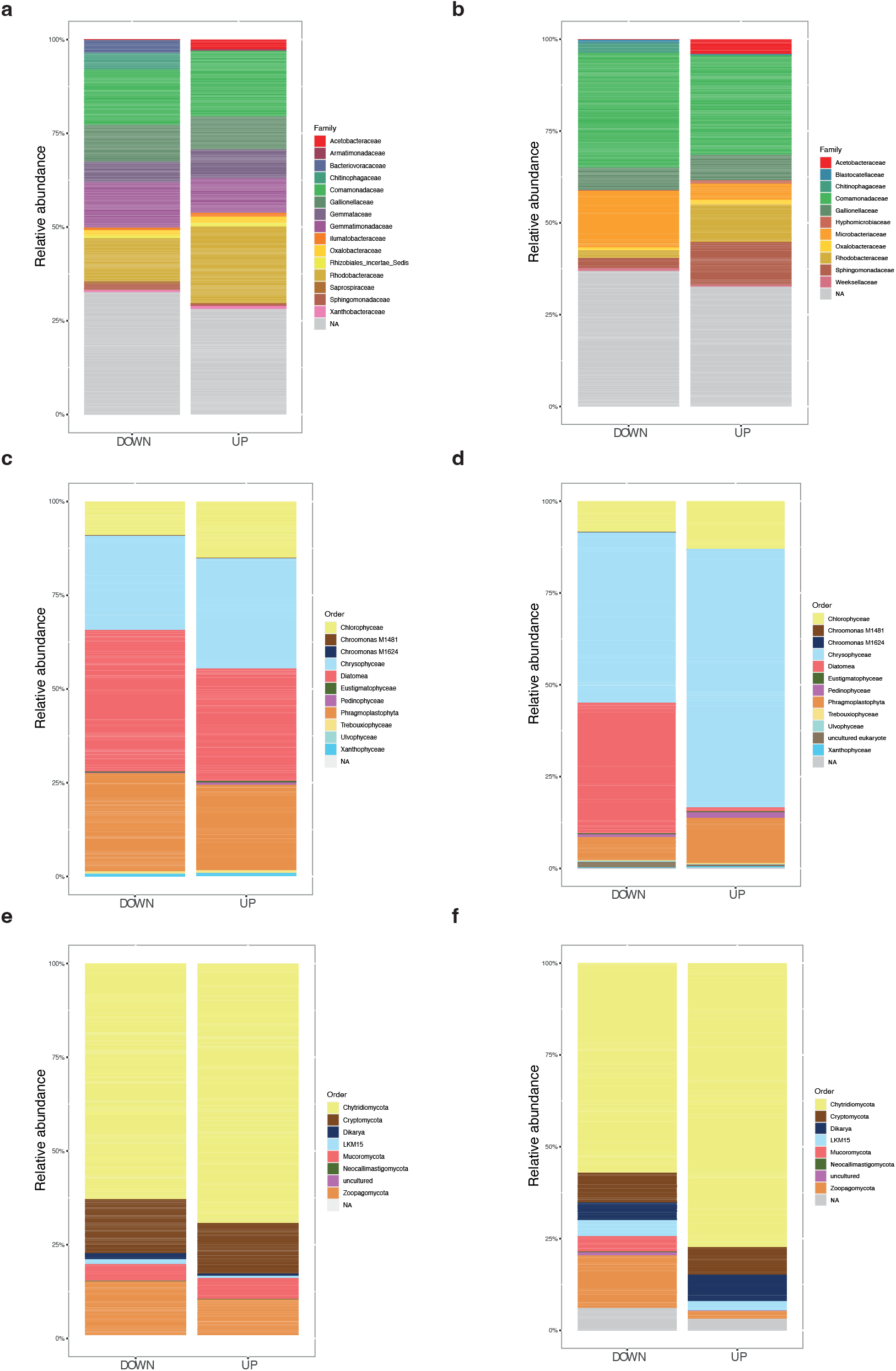
Taxa contributing to cross-domain interactions in Otemma. Relative abundance of prokaryotes found in the cross-domain networks of the (a) GFS and (b) TRIB in Otemma. (c) and (d) show the relative abundance of the phototrophs in the GFS and TRIB respectively, while (e) and (f) depict the relative abundance of the fungi in Otemma.

**Supplementary figure 3.**
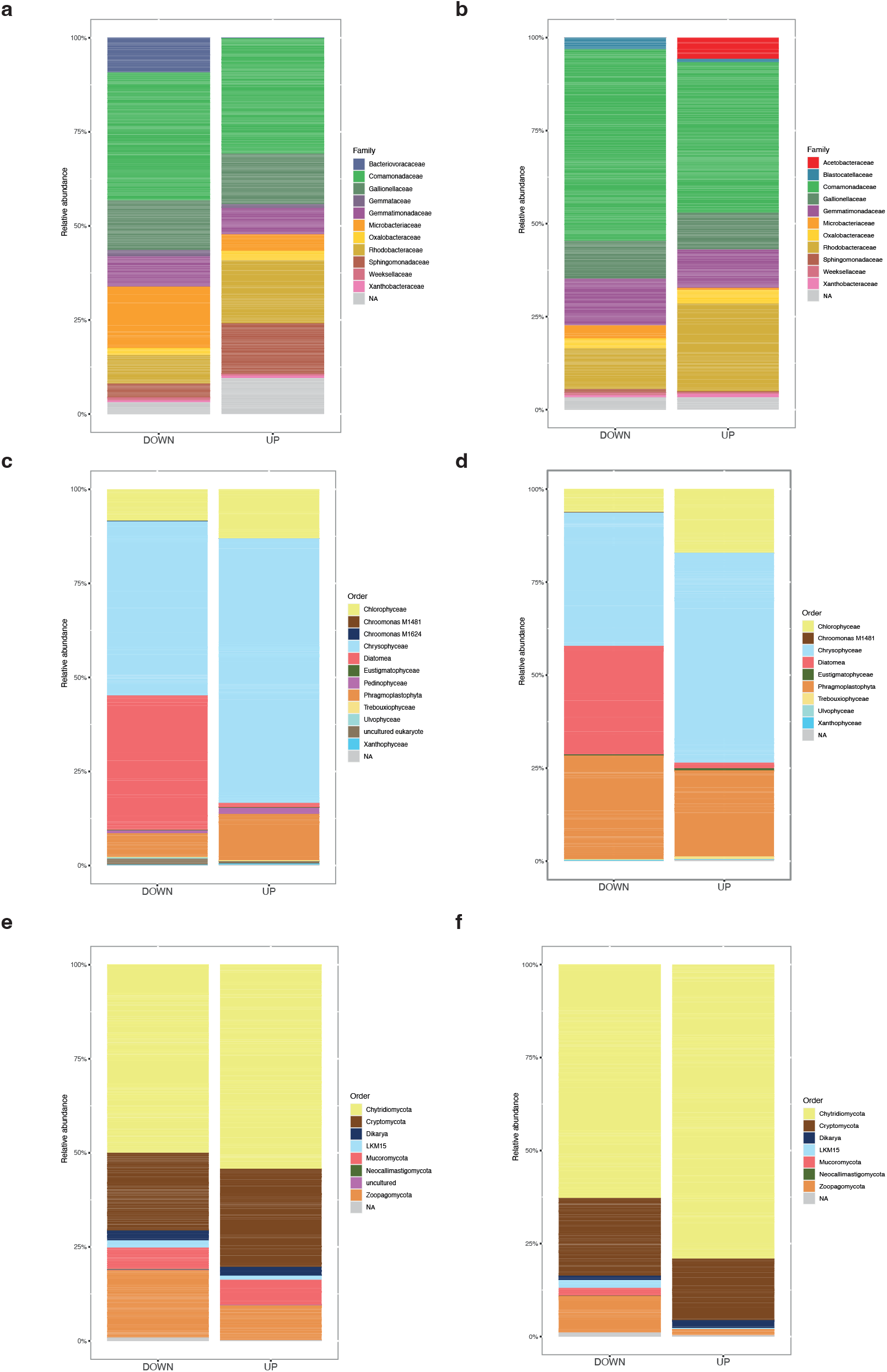
Taxa contributing to cross-domain interactions in Val Roseg. Relative abundance of prokaryotes found in Val Roseg in the cross-domain networks of the (a) GFS and (b) TRIB. Phototroph relative abundances in the (c) GFS and (d) TRIB. (e) and (f) depict the relative abundance of the fungi in GFS and TRIB in Val Roseg.

**Supplementary figure 4.**
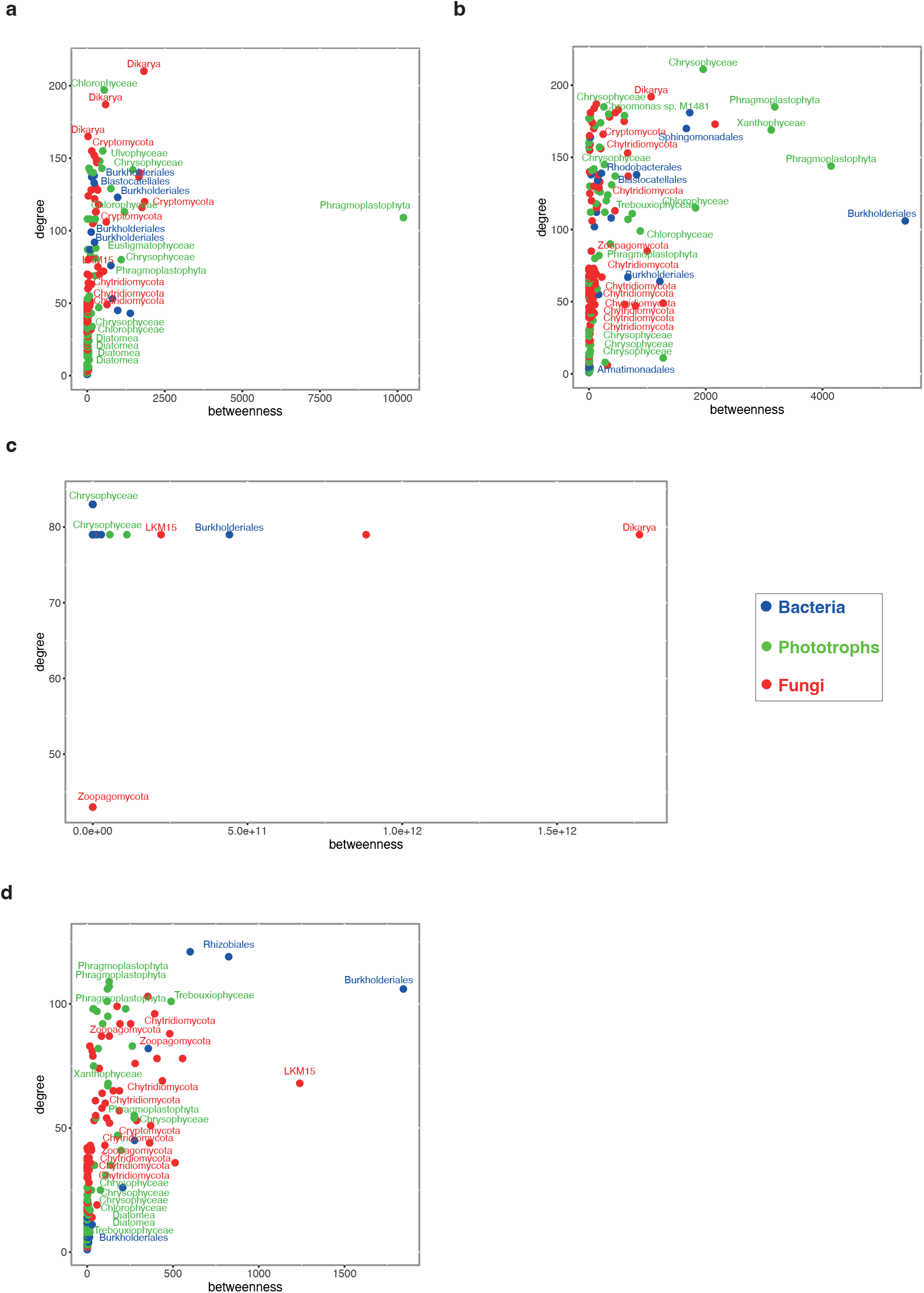
Keyplayer taxa in Otemma. The keyplayer taxa for the GFS at the (a) UP and (b) DOWN reaches are highlighted based on their domain of origin. Keyplayers in the TRIB at the (c) UP and (d) DOWN reaches from the TRIB are simultaneously shown. The x-axis represents the overall betweenness of the individual taxa, whereas the y-axis indicates the degree centrality.

**Supplementary figure 5.**
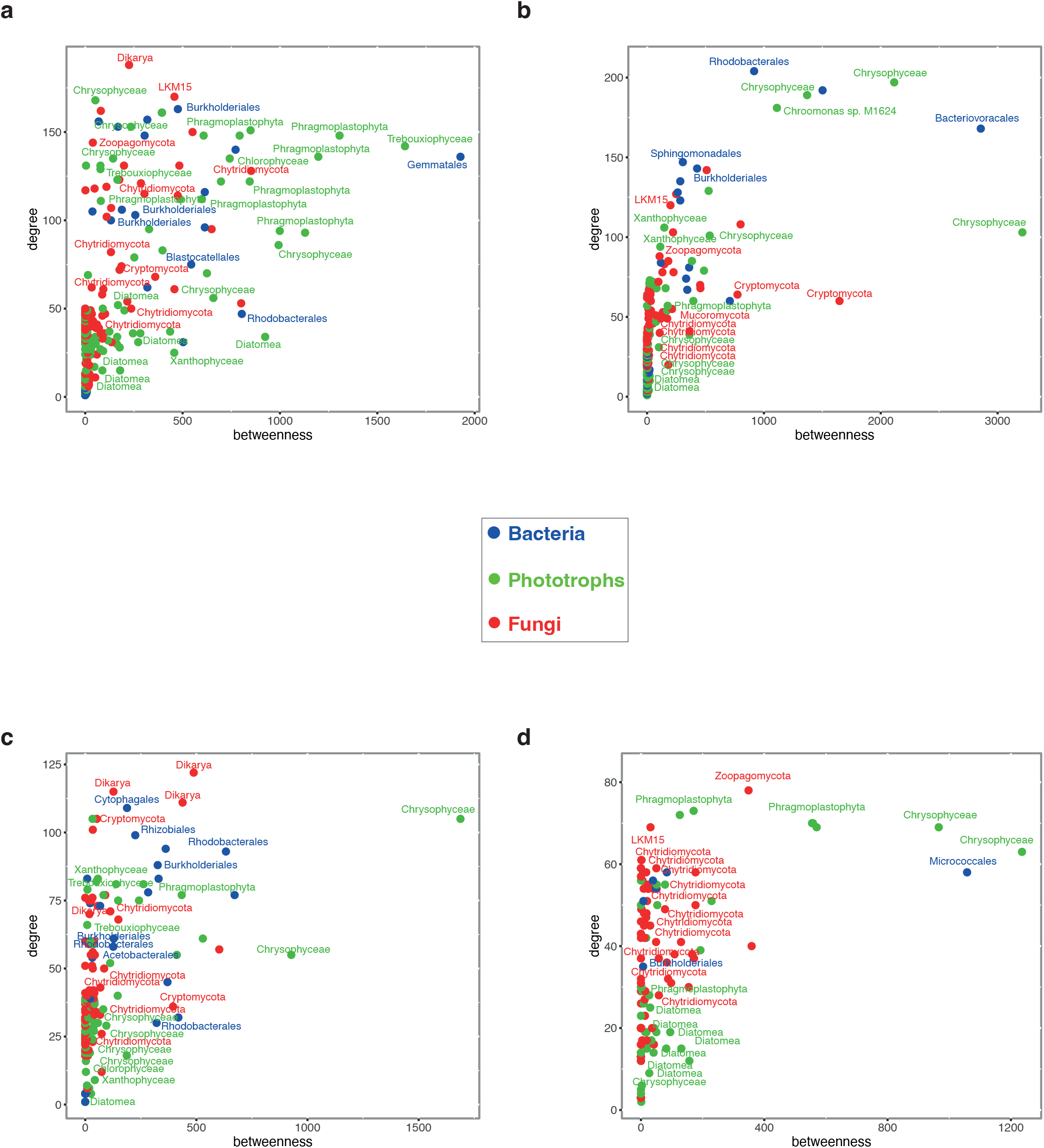
Keyplayer taxa in Val Roseg. The keyplayer taxa for the GFS at the (a) UP and (b) DOWN reaches in Val Roseg are highlighted. Keyplayers in the tributaries at the (c) UP and (d) DOWN reaches from the TRIB are depicted in the scatter plots. The x-axis represents the overall betweenness of the individual taxa, whereas the y-axis indicates the degree centrality, i.e., number of connections per node.

